# Outbreak Investigation of *Phytobacter diazotrophicus* Using Fourier Transform Infrared Spectroscopy compared to cgMLST

**DOI:** 10.1101/2025.09.17.676977

**Authors:** Débora Nicole de Oliveira Kulek, Geiziane Aparecida Gonçalves, Melise Chaves Silveira, Helena Regina Salomé D’Espindula, José Ferreira da Cunha Neto, Amanda Dal Lin, Thais Oliveira Gaio, Letícia Kraft, Ana Paula D’Alincourt Carvalho Assef, Marcelo Távora Mira, Marcelo Pillonetto

## Abstract

**Introduction:** *Phytobacter diazotrophicus* is an emerging opportunistic pathogen associated with hospital outbreaks and a reservoir of antimicrobial resistance genes; however, its clonal epidemiology remains poorly understood. Accurate identification and rapid clonal typing are crucial for infection control. Core genome MLST (cgMLST) is the genomic gold standard; however, it is both costly and time-consuming. Fourier-transform infrared (FT-IR) spectroscopy using the IR Biotyper^®^ emerges as a rapid and accessible typing alternative.

**Objective:** This study evaluated the performance of the IR Biotyper^®^ in discriminating *P. diazotrophicu*s outbreak isolates, compared to cgMLST.

**Methodology:** Eight epidemiologically relevant isolates, including two from a total parenteral nutrition (TPN) outbreak and four from a hemodialysis clinic, along with two outliers, were analyzed. Its identification was confirmed by API20E, MALDI-TOF MS, *nif*L gene qPCR, and WGS. Both cgMLST, based on 7,755 conserved genes, and the Fourier Transform Infrared (FT-IR) Spectroscopy by IR Biotyper^®^ were used for clonal typing, with the spectral cutoff empirically determined based on known clonal isolates.

**Results:** Both methodologies demonstrated remarkable agreement in detecting clonal clusters. The two TPN outbreak isolates showed 23 allele differences (ADs) by cgMLST and consistently clustered according to the IR Biotyper. Similarly, the four isolates from the outbreak at hemodialysis clinic formed homogeneous clusters by both methods (5-27 ADs by cgMLST). The outlier isolates were consistently discriminated.

**Conclusion:** FT-IR spectroscopy (IR Biotyper^®^) proved promising and complementary to cgMLST for typing *P. diazotrophicus* in outbreak scenarios, offering a rapid and cost-effective alternative for detecting clonal events. Its applicability in routine epidemiological surveillance is significant.

**Statement of Importance:** *Phytobacter diazotrophicus* is an emerging opportunistic pathogen of increasing concern in clinical environments, associated with hospital outbreaks and carrying antimicrobial resistance genes. Rapid detection and precise clonal characterization are essential for infection control and epidemiological surveillance. However, these are challenging for less-understood species. Genomic methods, such as cgMLST, are the gold standard, but their cost and time limitations hinder routine application in outbreak response. This study evaluated the IR Biotyper system, a rapid and accessible phenotypic tool that uses Fourier Transform Infrared Spectroscopy, compared to cgMLST for clonal typing of *P. diazotrophicus*. Our findings demonstrate a remarkable agreement between the two methods in identifying clonal clusters. This validation suggests the potential of the IR Biotyper as an effective tool for real-time screening and active surveillance of *P. diazotrophicus* in clinical settings. This enables a more agile response to outbreaks and aids in managing antimicrobial resistance, reserving high-resolution genomic analyses for more in-depth investigations.

## INTRODUCTION

The rapid and accurate identification of pathogens and the elucidation of their clonal relationships are fundamental pillars of clinical microbiology and public health laboratories. In hospital outbreak scenarios, the early detection of the responsible microorganism and the understanding of its dissemination dynamics are essential for implementing targeted control and prevention actions, minimizing the impact on morbidity and mortality (1–3). Atypical or emerging microorganisms, often underestimated in routine protocols, have gained relevance as etiological agents of outbreaks, especially those associated with the contamination of sterile solutions and medical devices (4–7).

In this context, *Phytobacter diazotrophicus* emerges as an opportunistic pathogen of growing concern in clinical environments (7). Although initially recognized for its environmental role, this Gram-negative bacillus has been increasingly linked to hospital outbreaks. For example, this bacterium was definitively associated with an outbreak of Total Parenteral Nutrition (TPN) contamination in Brazil in 2013 (5), and pediatric patients and hospital environment in Japan in 2023 (8). But many other previous outbreaks have been caused by this species but were wrongly assigned to *Pantoea* (*Enterobacter*) *agglomerans*, such as a nationwide outbreak in the 1970s in the USA (9,10) and in Brazil in 2010 (11). Also, another eight outbreaks were potentially caused by *P. diazotrophicus* but mistakenly attributed to *P. agglomerans* as reviewed (7). Unsurprisingly, its epidemiology and clonal dynamics are still insufficiently understood, partly due to its complex taxonomic evolution (7).

The precise identification of *Phytobacter diazotrophicus* has been a persistent challenge in clinical microbiology, often resulting in its misidentification by routine laboratory methods (7,12). Historically, this bacterium was classified within the *“Erwinia Herbicola-Enterobacter agglomerans”* complex (EEC) (13), which contributed to its confusion with other genera (12,14,15). *P. diazotrophicus* isolates are frequently misidentified as *Pantoea agglomerans, Kluyvera intermedia*, or *Leclercia adecarboxylata* by automated commercial systems and outdated identification databases (7). This difficulty largely stems from the high similarity of phenotypic identification, that is commonly used for routine species classification, resulting in misleading reference databases. There is also a similarity of the 16S rDNA gene sequences between these species, which limits the discriminatory capacity of this widely used molecular tool. Such misidentification directly compromises the epidemiological investigation of hospital outbreaks, impacts clinical and sanitary measures, and can affect the therapeutic approach, highlighting the importance of more efficient methodologies to ensure patient safety (6,7).

Futhermore, *P. diazotrophicus* has emerged as a significant reservoir of antimicrobial resistance (AMR) genes, among which we highlight some important carbapenemases (*bla*_KPC_, *bla*_NDM_, *bla*_IMP-6_, *bla*_GES-5_) and determinants associated with mobile genetic elements. Therefore, it underscores its potential for dissemination and its threat to therapeutic effectiveness (8,16–21). The rapid and precise identification of clonal lineages is, also essential for outbreak detection and epidemiological surveillance (2,22).

Alternatively, more laborious and time-consuming Whole Genome Sequencing (WGS) enables other analysis methods, such as core genome multilocus sequence typing (cgMLST), which is considered the gold standard for high-resolution typing, providing detailed insights into phylogenetic and epidemiological relationships (23)0). However, their large-scale application is often limited by high costs, long execution times, and the need for advanced bioinformatics infrastructure and expertise (1).

Given these limitations, Fourier transform infrared (FT-IR) spectroscopy has emerged as a promising alternative for bacterial typing. This phenotypic technique is fast, cost-effective, and more accessible, with promising results in discriminating strains of various clinically relevant species (1,24–26). The IR Biotyper^®^ (Bruker Daltonics, Bremen, Germany) is a commercial FT-IR platform that has been increasingly adopted in hospital microbiology laboratories due to its operational advantages (24,27).

However, the applicability and discriminatory power of FT-IR spectroscopy for the clonal analysis of rare or newly emerging species like *P. diazotrophicus* remain largely unexplored in the literature. Therefore, this study aims to evaluate the performance of FT-IR spectroscopy, using the IR Biotyper^®^ platform, in discriminating *Phytobacter diazotrophicus* isolates, compared to genomic typing based on cgMLST.

## MATERIALS AND METHODS

The experiments were conducted in a collaboration between three Brazilian institutions: the Center for Advanced Molecular Investigation of the Pontifícia Universidade Católica do Paraná (NIMA/PUCPR), the Central Laboratory of the State of Paraná (LACEN-PR), and the Laboratory of Bacteriology Applied to One Health and Antimicrobial Resistance (LabSUR) at the Oswaldo Cruz Institute (IOC/Fiocruz). The analyses were distributed among these laboratories according to the specificity of each methodological procedure.

### Ethical Considerations

The samples used in this study consisted exclusively of bacterial isolates, completely unlinked from any sensitive human patient information. Only epidemiological data and anonymized information, such as the institution of origin, collection date, and type of biological sample, were considered for the selection of isolates, ensuring privacy and compliance with data protection regulations. For research of this nature, national legislation waives the need for approval from the Research Ethics Committee (CEP).

## Description and Selection of Isolates

This study analyzed *Phytobacter diazotrophicus* isolates collected and identified from 2013 to 2021 and preserved in the strain bank of the LACEN-PR Bacteriology Sector at -80ºC. The initial identification of these isolates was performed according to the laboratory’s staged protocol, which includes API 20E^®^, Vitek-MS^®^ MALDI-TOF (Mazzetti et al, 2025 - in preparation), qPCR for *nif*L (28), and WGS. A total of eight isolates were used for this study to compare the methodologies. Among these, two isolates (5020RM and 5110RM) came from an outbreak that occurred in 2013, associated with the contamination of Total Parenteral Nutrition (TPN) by *P. diazotrophicus*, as previously described by Pillonetto, M. et al., 2018 (5). Additionally, four blood isolates (38394RM, 38397RM, 38453RM, and 38468RM), collected within five days at the end of 2021, were selected from a hemodialysis clinic (HDC), with a strong suspicion of representing an outbreak event. The two remaining isolates, 21916RM (rectal swab) and 32066RM (blood), from different periods, were included as outliers to evaluate the discriminatory capacity of the methodologies. Table 1 details the internal identification, origin, collection date, biological sample, and WGS identification for each of the eight strains used in the study.

**Table 1.**
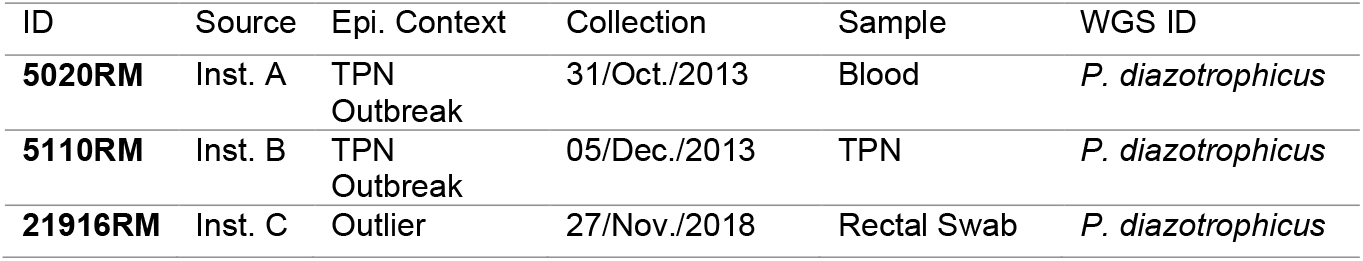
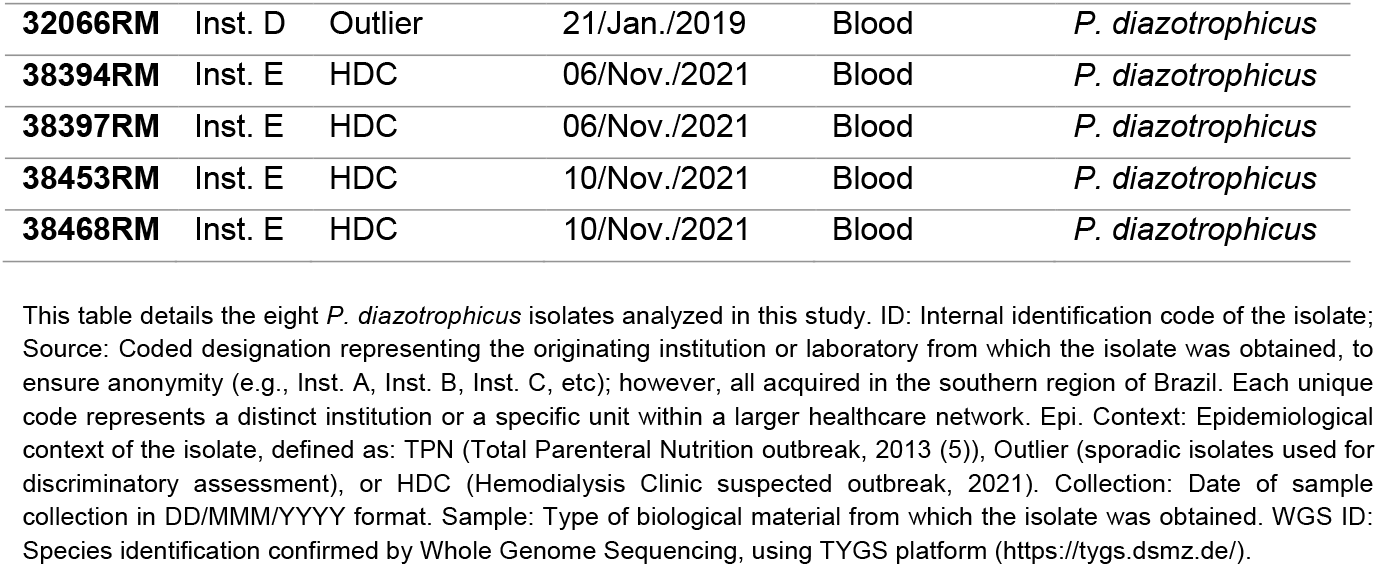
*P. diazotrophicus* isolates included in the study.

| ID | Source | Epi. Context | Collection | Sample | WGS ID |
| --- | --- | --- | --- | --- | --- |
| **5020RM** | Inst. A | TPN<br>Outbreak | 31/Oct./2013 | Blood | *P. diazotrophicus* |
| **5110RM** | Inst. B | TPN<br>Outbreak | 05/Dec./2013 | TPN | *P. diazotrophicus* |
| **21916RM** | Inst. C | Outlier | 27/Nov./2018 | Rectal Swab | *P. diazotrophicus* |
| **32066RM** | Inst. D | Outlier | 21/Jan./2019 | Blood | *P. diazotrophicus* |
| **38394RM** | Inst. E | HDC | 06/Nov./2021 | Blood | *P. diazotrophicus* |
| **38397RM** | Inst. E | HDC | 06/Nov./2021 | Blood | *P. diazotrophicus* |
| **38453RM** | Inst. E | HDC | 10/Nov./2021 | Blood | *P. diazotrophicus* |
| **38468RM** | Inst. E | HDC | 10/Nov./2021 | Blood | *P. diazotrophicus* |

This table details the eight *P. diazotrophicus* isolates analyzed in this study. ID: Internal identification code of the isolate; Source: Coded designation representing the originating institution or laboratory from which the isolate was obtained, to ensure anonymity (e.g., Inst. A, Inst. B, Inst. C, etc); however, all acquired in the southern region of Brazil. Each unique code represents a distinct institution or a specific unit within a larger healthcare network. Epi. Context: Epidemiological context of the isolate, defined as: TPN (Total Parenteral Nutrition outbreak, 2013 (5)), Outlier (sporadic isolates used for discriminatory assessment), or HDC (Hemodialysis Clinic suspected outbreak, 2021). Collection: Date of sample collection in DD/MMM/YYYY format. Sample: Type of biological material from which the isolate was obtained. WGS ID: Species identification confirmed by Whole Genome Sequencing, using TYGS platform (https://tygs.dsmz.de/).

## Microbiological and Molecular Characterization

### Culturing and Identification of Strains

Frozen strains were plated on MacConkey agar and incubated at 37 °C for 24 hours to confirm the viability and purity of the isolates. The identification of all strains was confirmed by API20E (bioMérieux, Marcy l’Étoile, France), MALDI-TOF MS (bioMérieux, Marcy l’Étoile, France), using the RUO (Research Use Only) Saramis® software (v. 4.0, bioMérieux) and a specific SuperSpectra® for the genus *Phytobacter* spp. This spectrum was built at LACEN-PR (Mazzetti et al., 2025, in preparation). Additionally, qPCR targeting the *nif*L gene (Nitrogen Fixation Regulatory Protein, one of the characteristics that originally define *Phytobacter* spp.) was performed in duplicate, according to a previously established protocol (28).

### Whole Genome Sequencing (WGS) and Initial Genomic Analyses

Genomic DNA for WGS was extracted with the PureLink^®^ Genomic DNA kit (Thermo Fisher Scientific, Carlsbad, CA, USA), and libraries were prepared with the TruSeq Nano DNA Library Preparation Kit for NeoPrep (Illumina, San Diego, CA, USA). Sequencing was performed on the Illumina MiSeq^®^ platform. The genomic analyses of the isolates were conducted using CABGen (29), and TYGS platform (https://tygs.dsmz.de/) (30). CABGen is a digital platform that integrates an automated bioinformatics pipeline for complete genomic analysis. It performs raw data quality control, genome assembly, and general quality analysis (considering parameters such as completeness, contamination, and size). Additionally, the tool generates genome annotation and sequence alignment for more in-depth analyses, such as the cgMLST schemes and phylogenetic tree construction. Detailed results, including contigs and contamination percentage, are available in Supplemental Table 01.

The precise species-level identification of *P. diazotrophicus* was additionally confirmed using the TYGS platform (30), which provided dDDH (digital DNA-DNA Hybridization) values (Supplemental Table 01) and the phylogenetic tree construction, with default parameters, using WGS.

### Clonality Analysis by cgMLST

Clonality analysis among *P. diazotrophicus* strains was conducted by core genome Multilocus Sequence Typing (cgMLST). The chewBBACA program (31) was used to create the cgMLST scheme and perform allele calling from the genomes. The cgMLST scheme was based exclusively on 13 *P. diazotrophicus* reference genomes available on NCBI (Supplemental Table 2), including clonal isolates from Japan deposited in GenBank(8). This scheme demonstrated the best discriminatory capacity and consistency for the species in the study, comprising 7,755 conserved genes (core genes). Allele identification and attribution were performed for each isolate, setting the execution mode to “1”, which in the chewBBACA program corresponds to the “allele calling” mode, a crucial step for translating genomic sequences into comparable allelic profiles. The number of allelic differences (AD) determined the genetic distance between the isolates. An allelic distance dendrogram was constructed using the Minimum Spanning Tree (MST) algorithm to visualize the clonal relationships between the strains submitting the allelic profiles to PHYLOViZ (https://online.phyloviz.net/index).

### Typing by Fourier-transform Infrared Spectroscopy - IR Biotyper®

Fourier transform infrared (FT-IR) spectroscopy is based on the absorption of infrared light by biomolecules (proteins, carbohydrates, and lipids) present in the bacterial cell. The analysis of the resulting spectra, which function as a “molecular fingerprint,” allows for the differentiation of microorganisms and the typing of closely related strains. The typing analysis of the eight isolates by FT-IR was performed using the IR Biotyper^®^ system (Bruker Daltonics), with OPUS V.8.2.28 software (Bruker Optics, Germany) and the IR Biotyper^®^ V4.0 software (Bruker Daltonics (32).

For this study, sample preparation, acquisition, and analysis of the spectra followed the manufacturer’s protocol (32). Briefly, pure bacterial colonies were grown on Mueller-Hinton agar (24 ± 0.5 hours at 37°C). Suspensions were prepared and applied in technical quintuplicate on dedicated silicon slides. Standard Bruker infrared test controls (IRTS 1 and IRTS 2) were pipetted in duplicate on each plate. This methodology was replicated in biological triplicate; however, the results obtained from these replicas did not demonstrate the necessary consistency to be included in the comparative analyses. The system’s default quality control was applied, excluding spectra that did not meet the predefined signal-to-noise ratio and interference criteria.

Spectral similarity analysis was performed with the processing conditions WN: 1300-800/cm^− 1^, Euclidean metric, and UPGMA (Unweighted Pair Group Method with Arithmetic Mean) linkage method. The software calculated the spectral distance between the spectra, allowing for the visualization of similarity between the samples in a dendrogram and 2D and 3D scatter plots using Linear Discriminant Analysis (LDA). The cutoff point (COV) for defining spectral similarity clusters was determined by a cross-validation approach, comparing the results with cgMLST and epidemiological data. This method was adopted due to the absence of a validated cutoff point for typing *P. diazotrophicus* by IR Biotyper, in line with the practices of previous studies (27,33–35).

The COV was precisely adjusted to ensure the spectral clusters reflected known biological and epidemiological relationships. This robust approach was achieved by calibrating the cutoff using two key sources: epidemiological data and cgMLST analysis. The known grouping of TPN outbreak isolates (5020RM and 5110RM) and the ability to discern epidemiologically unrelated strains (outliers) served as primary references. The results from parallel cgMLST analyses, a high-resolution genomic method considered the gold standard for clonal relationships, also served as a basis for evaluating the consistency of the groupings. This dual validation ensured that the IR Biotyper’s clustering was not based on an arbitrary mathematical value, but on strong evidence, guaranteeing that isolates with a confirmed clonal indication consistently clustered together in different epidemiological scenarios.

## RESULTS

### Culturing and Confirmation of Isolates

The eight selected isolates were consistently identified as *Pantoea* spp. 2 by API20E (Supplemental Table 3), as expected for *Phytobacter* species (7). They were consistently identified as *Phytobacter* sp. by MALDI-TOF MS Saramis, detecting the *nif*L, indicating confirmation of the genus *Phytobacter* spp. (Supplemental Tables 4 and 5, respectively) and as *Phytobacter diazotrophicus* by WGS (Supplemental Table 1). The whole genome-based phylogenetic analysis (Supplemental Figure 1) confirmed the species-level identification, demonstrating that all isolates cluster robustly with the reference strain *P. diazotrophicus* DSM 17806. The tree topology also corroborates the clonality findings, with isolates from TPN outbreak ant the hemodialysis clinic forming distinct clades.

### Clonality Analysis by cgMLST

The results of cgMLST typing are presented in the dendrogram in Figure 1. The analysis revealed distinct clonal clusters among the *P. diazotrophicus* isolates. Isolates 5020RM and 5110RM, previously associated with the 2013 TPN outbreak (5), showed 23 allelic differences and were consistently grouped (See Figure 1, Cluster A). Similarly, the four blood isolates (38394RM, 38397RM, 38453RM, and 38468RM) from the same hemodialysis clinic, collected in 2021, formed a genotypically homogeneous cluster, as shown in Figure 1 (Cluster B). The difference between these isolates ranged from 5 to 27 AD, with 38394RM being central to the cluster (5 AD from 38468RM, 6 AD from 38453RM, and 27 AD from 38397RM). The other isolates showed greater genetic diversity and were distributed as ungrouped isolates in the complete dendrogram.

**Figure 1.**
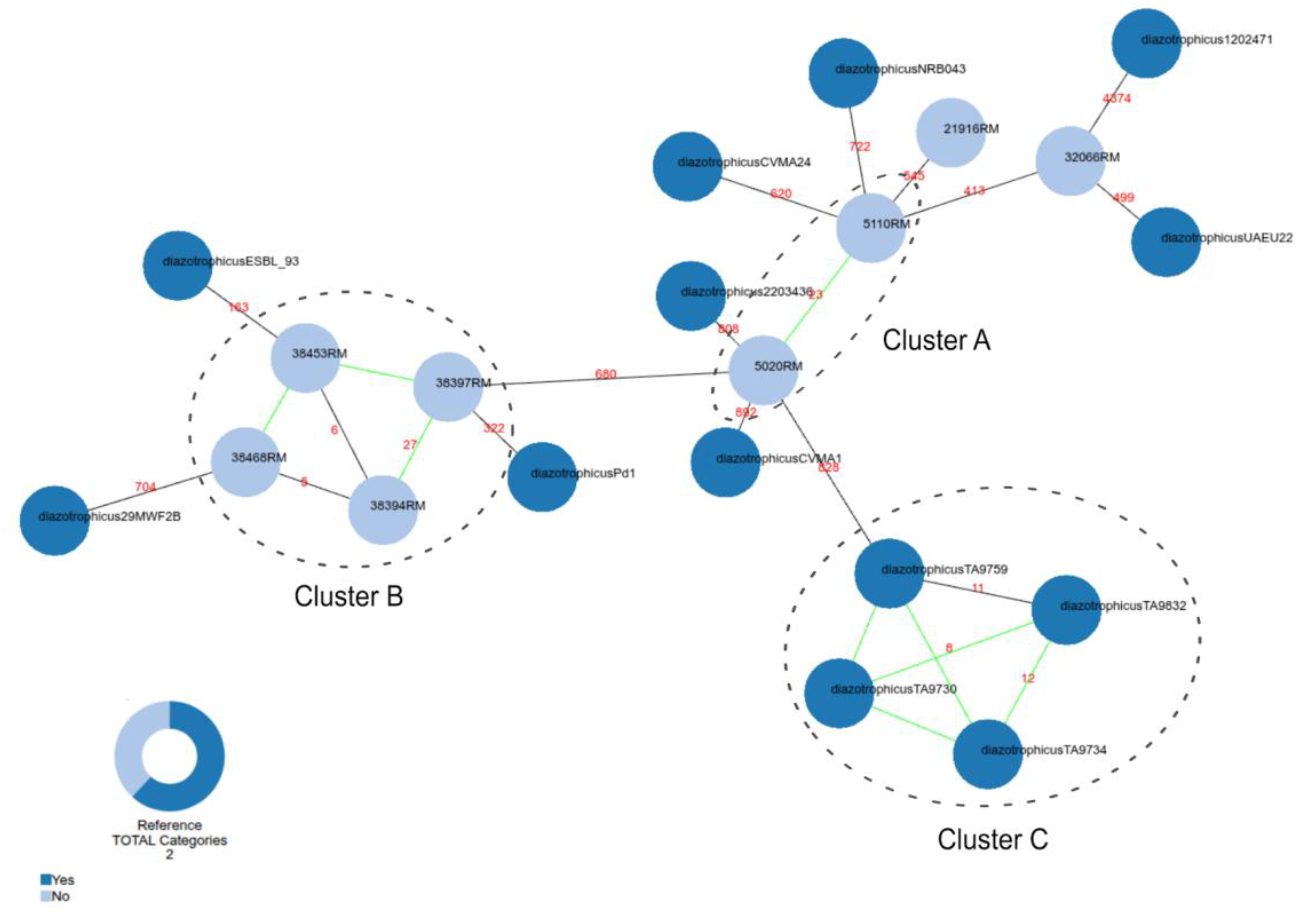
Allelic distance dendrogram of *P. diazotrophicus* by cgMLST (7,755 core genome genes). The dendrogram illustrates the clonality relationships between the analyzed *P. diazotrophicus* isolates and reference strains from NCBI. The connecting lines represent the genetic relationships, and the numbers in red indicate the allelic differences (AD). Green connection lines represent connections with allelic differences equal to or lower than 12. Black circles with dashed linehighlight the main clonal clusters identified: Cluster A corresponds to the 2013 TPN outbreak, which includes isolates 5020RM and 5110RM; Cluster B indicates the cluster associated with the hemodialysis clinic, composed of isolates 38394RM, 38397RM, 38453RM, and 38468RM, and Cluster C, representing four clonal isolates from Japan (8), already deposited at geneBank. Isolates that are not within these circles represent strains without a direct epidemiological correlation and with a lower degree of clonal relationship within the analyzed set.

### Typing Analysis by Fourier-transform Infrared Spectroscopy - IR Biotyper®

For all eight strains, spectra were produced in quintuplicate. One of the spectra of strain 38397RM was excluded from the analysis because it showed a discrepant profile in relation to the other replicates, compromising the proper formation of the grouping. The spectral similarity analysis was performed with the cutoff calibrated to a value of 0.203, in accordance with the cross-validation approach described in the methodology, using both cgMLST and epidemiological data. Applying this cutoff ensured that the spectral relationships accurately reflected the confirmed clonal groupings. The cophenetic correlation coefficient was 0.980. The results of the spectral similarity relationships are presented in a hierarchical clustering dendrogram (Figure 2), complemented by 2D and 3D scatter plots (Figures 3 and 4).

**Figure 2.**
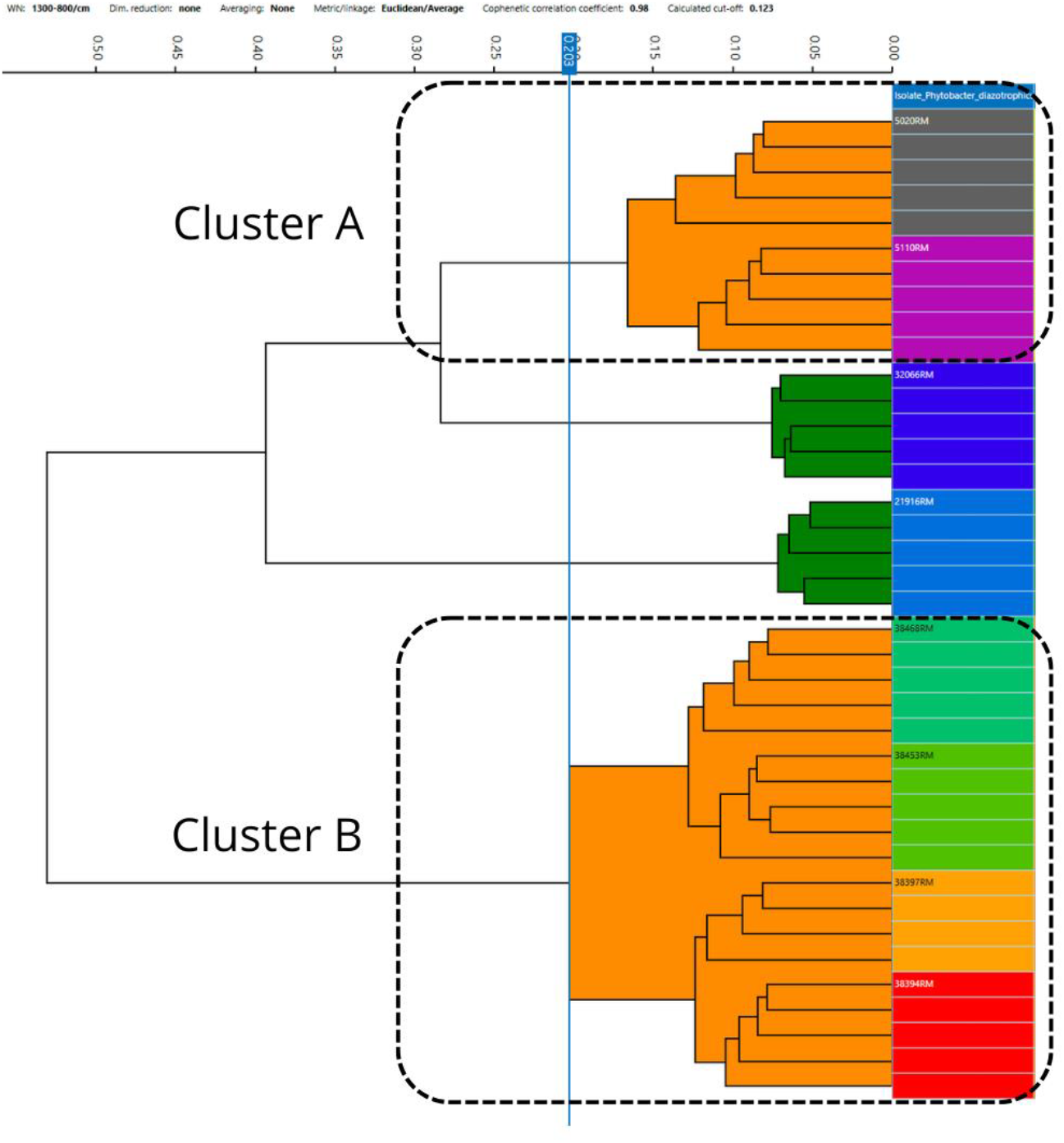
Hierarchical clustering dendrogram of *Phytobacter diazotrophicus* isolates by IR Biotyper®. The dendrogram illustrates the spectral similarity relationships between *P. diazotrophicus* isolates through hierarchical clustering. The analysis was performed based on Euclidean distance and the UPGMA linkage method. The dendrogram demonstrates the formation of clusters consistent with the epidemiological events (Cluster A - 2013 TPN outbreak, and Cluster B - associated with the hemodialysis clinic). The high consistency of the technical replicates is visually represented by the colors of the branches, with different colors indicating distinct isolates or clusters. The orange groups represent the clonal isolates with a confirmed epidemiological correlation (COV 0.203), while the green groups contain the isolates that were outliers without a direct epidemiological link to the others. The clustering reflects the validated clonal relationships, where isolates from the TPN outbreak and the hemodialysis clinic form cohesive groups, while the unrelated isolates (outliers) appear on separate branches.

**Figure 3.**
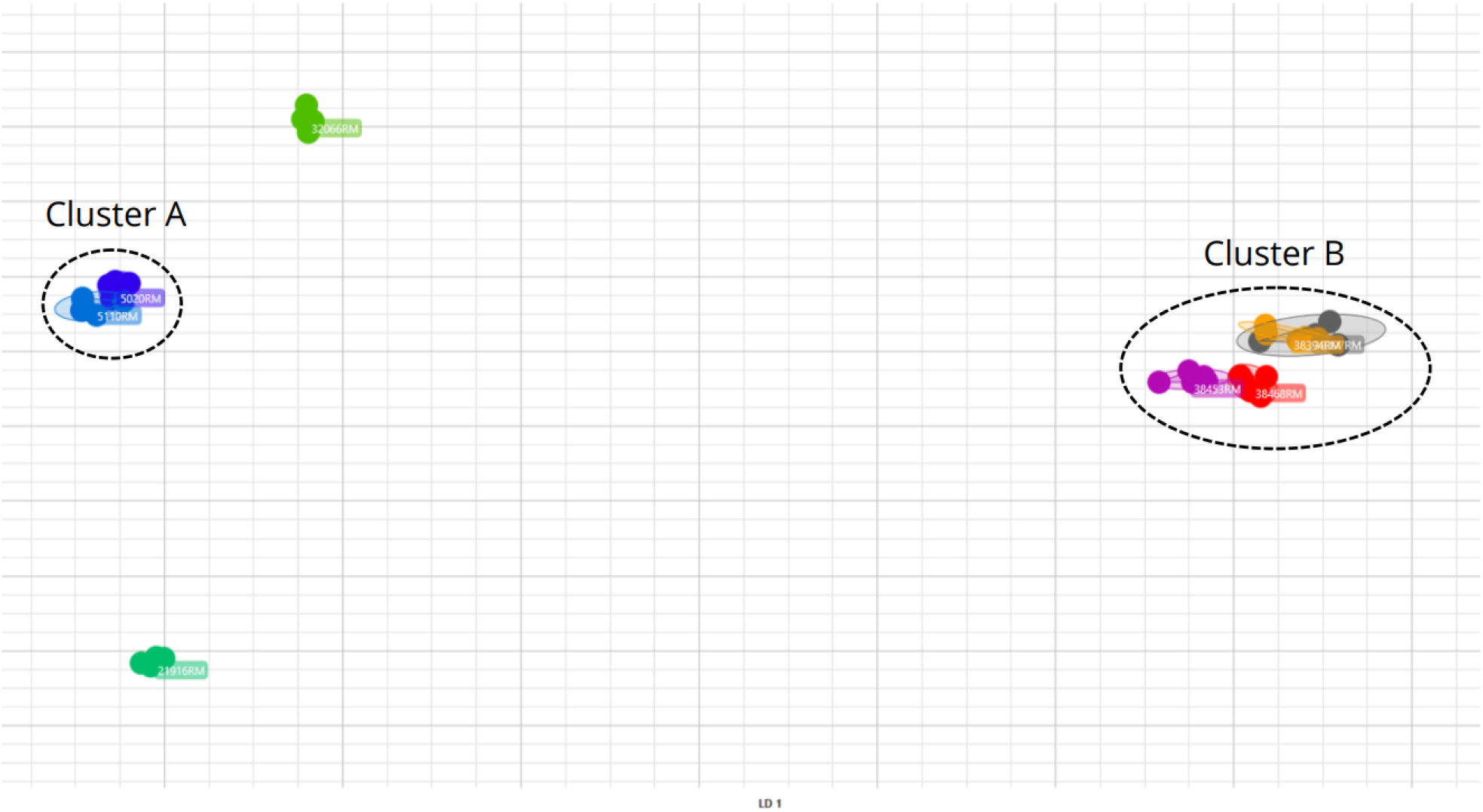
2D Scatter Plot by LDA of the spectral similarity relationships of *Phytobacter diazotrophocus* isolates by IR Biotyper® (1300-800 cm^− 1^). The 2D scatter plot (LDA, with dimensionality reduction of 5 Principal Components (PCs), explaining 96.2% of the variance) illustrates the spectral similarity between the eight *P. diazotrophicus* isolates analyzed by the IR Biotyper® system. Each point represents a technical replicate of an isolate, and the isolates are grouped by color and labeled with their respective IDs to indicate epidemiological relationships. Cluster A (TPN): Isolates 5020RM (dark blue) and 5110RM (light blue) group into a cohesive cluster, separate from the other groups, reflecting their known clonality. Cluster B (Hemodialysis Clinic): Isolates 38397RM (gray), 38394RM (orange), 38453RM (magenta), and 38468RM (red) form a compact and close cluster, indicating a clonal event. Outlier Isolates: Isolates 21916RM and 32066RM, identified in distinct shades of green, are clearly distant from the main clusters and from each other

**Figure 4.**
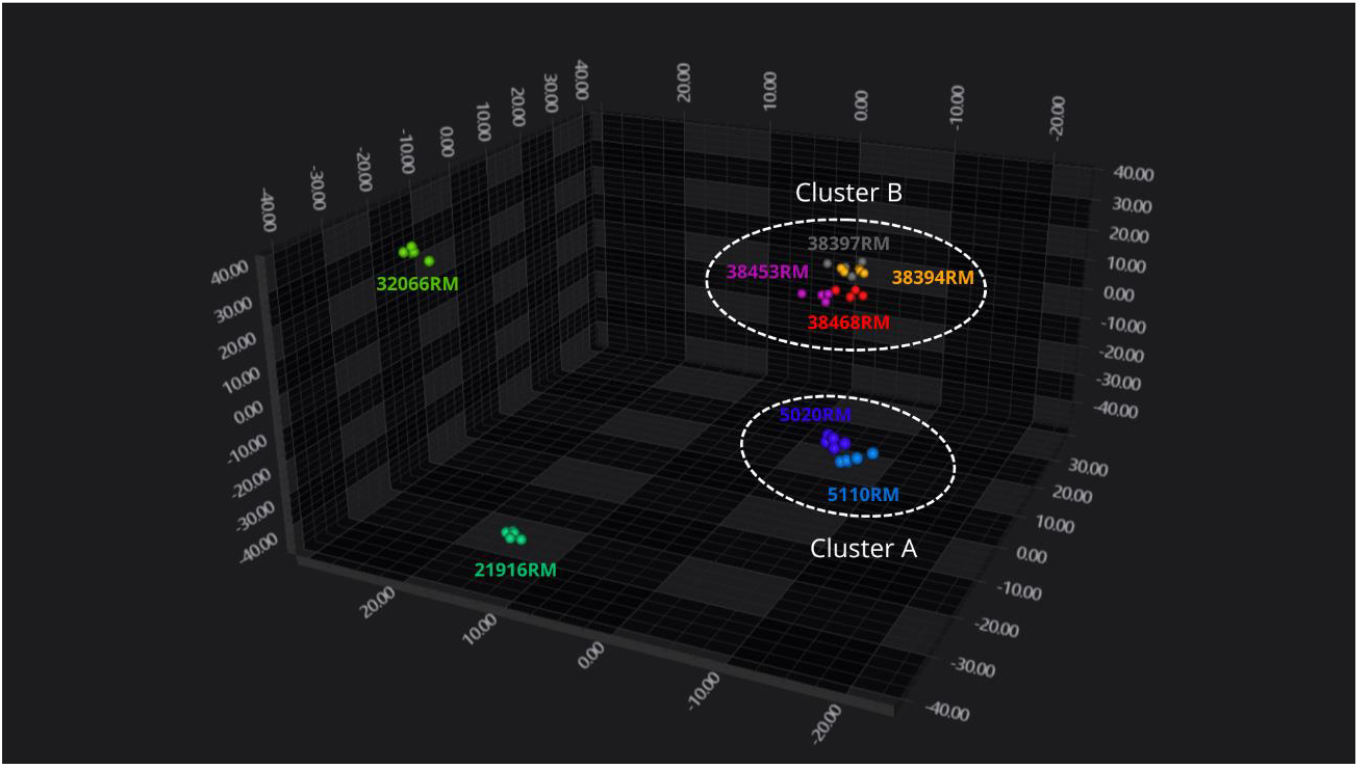
3D Scatter Plot of the spectral similarity relationships of *Phytobacter diazotrophicus* isolates by IR Biotyper® (1300-800 cm^− 1^). The 3D scatter plot (LDA, with dimensionality reduction of 10 PCs explaining 96.2% of variance) illustrates the spectral similarity between the eight *P. diazotrophicus* isolates analyzed by the IR Biotyper® system. Each point represents a technical replicate of an isolate, and the isolates are grouped by color and labeled with their respective IDs to indicate epidemiological relationships. Cluster A (TPN): Isolates 5020RM (dark blue) and 5110RM (light blue) groups into a cohesive cluster, separated from the other groups, reflecting their known clonality. Cluster B (Hemodialysis Clinic): Isolates 38394RM (orange), 38397RM (gray), 38453RM (magenta), and 38468RM (red) form a compact and close cluster, indicating a clonal event. Outlier Isolates: Isolates 21916RM and 32066RM (both in distinct shades of green) are distant from the main clusters and from each other, demonstrating the discriminatory capacity of the method to separate unrelated strains. The proximity of the points indicates high spectral similarity, while the distance suggests dissimilarity.

The dendrogram generated by the IR Biotyper^®^ demonstrated the formation of clusters consistent with epidemiological events. The isolates from the 2013 Total Parenteral Nutrition (TPN) outbreak, 5020RM and 5110RM, grouped strongly, forming a cohesive cluster. Similarly, the four isolates from the hemodialysis clinic (38394RM, 38397RM, 38453RM, and 38468RM) also formed a compact and well-defined cluster, revealing high spectral similarity among themselves. Analysis of the technical replicates showed high coherence, indicating a consistent grouping of the replicates of the same isolate. The isolates 21916RM and 32066RM, strains without a direct epidemiological correlation with the others (outliers), were distinct from the main clusters and each other, located on separate branches of the dendrogram. This separation demonstrates the IR Biotyper^®^’s ability to discriminate between related and unrelated strains, based on Euclidean distance and the UPGMA method. The 2D and 3D scatter plots (Figures 3 and 4, respectively) corroborated these observations, grouping the isolates into clusters consistent with the epidemiological events by Linear Discriminant Analysis (LDA).

## DISCUSSION

*P. diazotrophicus* isolates with clinical epidemiological relatedness were used to compare the performance of core genome MLST (cgMLST) and Fourier transform infrared spectroscopy (IR Biotyper®) typing to elucidate clonal events for this species. Clonality analysis by cgMLST (Figure 1) demonstrated high discriminatory capacity and consistent grouping of the isolates, corroborating the epidemiological data. The distinction between an outbreak, defined epidemiologically by the occurrence of cases in a specific location and period, and clonality, which refers to the genetic relationship between microorganisms, is fundamental for interpreting these findings. Genomic methods, such as cgMLST, are molecular tools to confirm whether a genetically related bacterial lineage causes an epidemiological outbreak event. There is no publicly available or established cgMLST scheme specifically for *P. diazotrophicus*, so we have created one using high-quality genomes available at NCBI in August 2025.

The whole genome-based phylogenetic tree topology (Supplemental Figure 1) provided a robust basis for species-level identification and corroborated the clonality findings by cgMLST. The phylogenetic analysis revealed a critical point of taxonomic confusion, as genomes previously deposited as *Citrobacter bitternis* JCM 300009 and *Kluyvera intestini* GT-1 clustered robustly within the *Phytobacter diazotrophicus* clade. This observation is corroborated by a recent genomic study that formally reclassifies these taxa as heterotypic synonyms of *P. diazotrophicus*, based on comprehensive comparative analyses including 16S rRNA gene phylogeny, core genome phylogeny, ANI, and dDDH (36). This compelling evidence highlights the historical misidentification of this genus and underscores the need for modern genomic approaches to ensure accurate species identification.

In this sense, isolates 5020RM and 5110RM, from the 2013 TPN outbreak, showed 23 allelic differences (AD), confirming the expected clonality, previously determined by Pillonetto et al, 2018a (5), using repPCR. Similarly, the four isolates from the hemodialysis clinic (38394RM, 38397RM, 38453RM, and 38468RM) formed a genotypically homogeneous cluster, with ADs ranging from 5 to 27. These findings, when interpreted in the context of the high genomic plasticity of *Phytobacter* (37), suggest that we are facing a continuous transmission event where the microevolution of the original strain resulted in genetic and phenotypic variants. Isolate 38397RM, for example, showed a phenotypic difference (positivity for melibiose, as per Supplemental Table 3) and a higher number of ADs (up to 27 ADs) than other cluster B members.

The cutoff point of 27 AD to define clonality in *P. diazotrophicus* was adopted based on epidemiological contexts and literature for species with limited population data and the absence of a validated threshold in the literature (38,39). Although a threshold of ≤10 AD is often used (40–42), a criterion of >10 AD has been established in previous studies for poorly described species and also based on epidemiological contexts (38,39). Therefore, the congruence between epidemiological and genomic data (even with higher ADs) and the known plasticity of the species is the basis for confirming that the HDC isolates do represent a clonal outbreak event.

In parallel, typing by IR Biotyper® proved to be a promising tool for evaluating clonality in *P. diazotrophicus*. The dendrogram (Figure 2) and the 2D and 3D scatter plots (Figures 3 and 4) revealed a clear separation of the isolates into well-defined clusters, which align perfectly with the epidemiological and genomic data. The isolates from the TPN outbreak and the hemodialysis clinic formed separate cohesive clusters, while the outliers (21916RM and 32066RM) were consistently distinct. The absence of a previously established spectral similarity cutoff for *P. diazotrophicus* by IR Biotyper® represents a common challenge for applying new technologies in emerging or poorly characterized species.

Our study addressed this gap through an empirical determination of the cutoff point, a crucial strategy for initially validating the technique in outbreak contexts. By using as a reference both the isolates with known confirmed clonality from the TPN outbreak and the suspected grouping of the isolates from the hemodialysis clinic (whose strong clonal indication was confirmed by parallel cgMLST analyses), we were able to establish a spectral similarity threshold (0.203) that demonstrated a consistent correlation with genomic clonality. This pragmatic approach allowed for the interpretation of IR Biotyper^®^ groupings for *P. diazotrophicus* and illustrates a viable model for the validation and use of the technique in other microorganisms with limited population data, facilitating the translation to routine laboratory work. The ability of the IR Biotyper^®^ to discern clonal isolates with this empirically determined cutoff reinforces its potential as a valuable tool for outbreak surveillance (34,35).

The comparison between cgMLST and IR Biotyper® revealed a remarkable agreement in identifying clonal clusters. Both methods were able to discriminate genetically distant isolates and successfully group outbreak-related strains, confirming clonality in two distinct events. cgMLST, a methodology based on WGS, offers the highest molecular resolution and allows for the precise quantification of genetic differences (ADs), which are necessary for in-depth epidemiological investigations and evolutionary studies. On the other hand, the IR Biotyper^®^ stands out for its speed, operational simplicity, and low cost per sample, presenting itself as a valuable screening tool. Its ability to provide typing information in approximately four hours, with a level of discrimination comparable to cgMLST for the detection of outbreak clusters in *P. diazotrophicus*, suggests its potential for routine, real-time epidemiological surveillance and for screening the selection of samples for more expensive and detailed WGS analyses.

Limitations of the study include the restricted number of isolates analyzed, which does not allow for the definition of robust population patterns for *P. diazotrophicus* in different geographical regions. This is a limitation inherent to the still large unknowingness of its pathogenic role and its difficulty in identification. Additionally, the empirical determination of cutoff points for clonality, although justified by the absence of data in the literature for this emerging species, requires validation by multicenter studies and with greater genetic diversity. The inconsistency observed in some of the results of the biological replicates of the IR Biotyper® also represents a limitation, suggesting the need to optimize preparation protocols or spectrum acquisition to ensure reproducibility in future evaluations.

## CONCLUSIONS

This study demonstrates that FT-IR spectroscopy is a promising and alternative tool to cgMLST for the screening and typing of *P. diazotrophicus* in outbreak scenarios, offering a rapid and cost-effective alternative for detecting clonal events. The continuous surveillance of this species, combined with the strategic application of both methodologies, is fundamental for understanding its epidemiology and for developing effective strategies for controlling healthcare-associated infections.

